# Using artificial intelligence on dermatology conditions in Uganda: A case for diversity in training data sets for machine learning

**DOI:** 10.1101/826057

**Authors:** Louis Henry Kamulegeya, Mark Okello, John Mark Bwanika, Davis Musinguzi, William Lubega, Davis Rusoke, Faith Nassiwa, Alexander Börve

## Abstract

**Introduction:** Artificial intelligence (AI) in healthcare has gained momentum with advances in affordable technology that has potential to help in diagnostics, predictive healthcare and personalized medicine. In pursuit of applying universal non-biased AI in healthcare, it is essential that data from different settings (gender, age and ethnicity) is represented. We present findings from beta-testing an AI-powered dermatological algorithm called Skin Image Search, by online dermatology company First Derm on Fitzpatrick 6 skin type (dark skin) dermatological conditions.

**Methods:** 123 dermatological images selected from a total of 173 images retrospectively extracted from the electronic database of a Ugandan telehealth company, The Medical Concierge Group (TMCG) after getting their consent. Details of age, gender and dermatological clinical diagnosis were analyzed using R on R studio software to assess the diagnostic accuracy of the AI app along disease diagnosis and body part. Predictability levels of the AI app was graded on a scale of 0 to 5, where 0-no prediction made and 1-5 demonstrating reducing correct prediction.

**Results:** 76 (62%) of the dermatological images were from females and 47 (38%) from males. The 5 most reported body parts were; genitals (20%), trunk (20%), lower limb (14.6%), face (12%) and upper limb (12%) with the AI app predicting a diagnosis in 62% of image body parts uploaded. Overall diagnostic accuracy of the AI app was low at 17% (21 out of 123 predictable images) with varying predictability levels correctness i.e. 1-8.9%, 2-2.4%, 3-2.4%, 4-1.6%, 5-1.6% with performance along individual diagnosis highest with dermatitis (80%).

**Conclusion:** There is a need for diversity in the image datasets used when training dermatology algorithms for AI applications in clinical decision support as a means to increase accuracy and thus offer correct treatment across skin types and geographies.

## Introduction

Dermatological conditions continue to be among the leading cause of non-fatal disease burden globally despite not receiving the required attention in terms of prevention, policy and clinical management [1,2]. When comparing absolute years lived with disability (YLDs), the global burden of disease study reported skin diseases to be the fourth leading cause of disability, after iron-deficiency anemia, tuberculosis, and sense organ diseases [3]. The prevalence rates of skin diseases appear to take on a geographical and socio-economic variation where limited resources settings like in sub-Saharan African, high prevalence of dermatological conditions like viral warts, pyoderma, cellulitis, scabies, psoriasis, alopecia areata, urticaria, fungal skin diseases, and decubitus ulcers have been reported [4].

Little research has been documented on the prevalence and burden of skin diseases in Uganda. Studies show that a number of dermatological conditions are endemic in some regions of the country, for example, *Visceral Leishmaniasis* is endemic in the north-eastern districts of Moroto and Kotido [5]. Trachoma is endemic in the northern districts whereas Buruli ulcer in the western and north-western parts of the country. A few other skin diseases like leprosy and guinea worm have been controlled despite pockets continue to persist in some areas. Studies on neglected tropical diseases in Uganda show that soil-transmitted helminths, schistosomiasis, lymphatic filariasis, and onchocerciasis all being dermatological diseases have a big socioeconomic impact on the affected subpopulations in Uganda with the related reduced productivity from disease state and cost of treatment [6].

A number of factors contribute to this observed trend in the occurrence of skin diseases in Uganda and these include; living conditions that bring humans closer to the vector e.g. brick and mud huts, close proximity with water bodies, poor disposal of human waste and sharing homesteads with animals among others. These conditions provide a favorable environment for the proliferation of skin diseases like skin fungus and embryonic blood flukes that nestle into their human hosts [7]. These situations are further boosted by water insecurity in the communities coupled with the lack of proper treatment support systems in our limited resource settings [8].

In a country where the doctor to patient ratio stands at an approximate 1:25,000 [9], and worse for specialist’s like dermatologist where cost of consultation is about 12 USD [10] access to health professionals more so specialist is very challenging. This means that for health conditions that would require a specialist opinion often go undiagnosed and untreated causing personal social issues and hardships on their families with associated morbidity. Innovations like the use of Artificial Intelligence (AI) aided platforms as a diagnostic tool to assist in diagnosing and choosing the correct treatment plan on the first visit could help to bridge the gap of low numbers of dermatologists in limited resource settings.

Techopedia Inc defines AI “as an area of computer science that emphasizes the creation of intelligent machines that work and react like humans” [11]. The use of AI has already shown immense potential with its integration in industries like the automobile in driverless cars, financial markets, language translation among others [12]. This has not left the health sector with many AI based innovations successfully being applied for image analysis in radiology, pathology, and dermatology with its benefits of dramatically reducing the cost of medical care while providing quicker diagnosis [13].

Concerns about the diagnostic accuracy of deep learning computer vision aided tools in healthcare have been noted, a study analyzing the performance of an imaging analysis computer algorithm showed tendencies of overdiagnosis [14]. A number of factors have been attributed to this suboptimal performance of computer vision tools in health and among them is the lack of diversity in datasets used for training the algorithms which often excludes certain demographics especially blacks [15,16].

New AI models in dermatology are continuously developed as more image databases are available and computer power is relatively cheap. Every year the International Skin **Imaging** Collaboration (ISIC) organize a challenge with the goal to develop image analysis tools to enable the automated diagnosis of melanoma from dermoscopic images [xx - https://challenge2018.isic-archive.com/]. The limitations of the accuracy and diversity of the models are images. Thus models will continuously get better and my diverse with time, when these images are collected and available.

In the majority of scenarios, these AI-based solutions are designed from the western and European world with their algorithms trained with western/European world environments. This by default leaves out specific demographics who in most cases these solutions are being designed to solve their solutions; a fallacy! It is important to note that machine learning algorithms sift through millions of training datasets/data points to make correlations and predictions, as such developing models that embrace people from different backgrounds and communities is critical to having an all-inclusive AI experience. In view of this, we carried out a study to assess the diagnostic accuracy of an AI algorithm developed by First Derm using a deep convolutional neural network (CNN) called “Skin Image Search” [17] on Fitzpatrick 6 skin types (black-dark) dermatological conditions.

## Materials and Methods

### Study Design

An observational study with descriptive statistical analysis of independent datasets containing Fitzpatrick 6 skin type dermatological conditions that were retrospectively extracted from the Telehealth platforms of a digital health company called The Medical Concierge Group (TMCG), located in Kampala-Uganda [18].

### Data Collection

TMCG operates a 24/7 digital health platform manned by qualified and licensed doctors that provide remote resolution to users’ medical inquiries. Dermatological inquiries involve users sharing images of the skin condition via the tele-platforms using their smartphones with additional history taking by the clinician to assess associated factors including; onset, pattern, exacerbating or relieving factors, chronicity among others as means to reach the correct diagnosis. Each individual image was reviewed by 3 independent general practitioners for which the final clinical diagnosis of the image was the most reported diagnosis. In instances where there was a disagreement, the final diagnosis was achieved by consensus. The shared images were anonymized prior to storage in the electronic medical records database. In this study, dermatological images between January to March 2018 were selected according to inclusion and exclusion criteria (see below). Data collection was done by consecutive sampling, where images were collected until the desired sample size i.e. dermatological images shared from January to March 2018 was acquired. All included images were imported by their file names into Microsoft Excel 2019 [19] and the following additional information was added; patient ID, gender, body site and final diagnosis (defined as the clinical diagnosis).

### Inclusion Criteria

All consecutive dermatological images within the tele-platform at TMCG were included. All images were anonymous since they contained no identifying data and the patient could not be recognized by the image. When necessary this was achieved by cropping the image in Microsoft paint [19].

### Exclusion Criteria

Non accessible images due to image quality, (for example bad lighting or blurred focus) or unfit image composition (camera not aimed at skin lesion, image taken from inadequate distance or angle) were excluded. Images showing pen markings like circles or arrows were excluded as well as where the skin lesions were wholly or partially covered by a dressing e.g. bandage. Images with more than one diagnosis were cropped in a way that the confirmed diagnosis became the main visible lesion in the image.

Skin Image Search™ – an artificial intelligence Dermatology and Venereology classification system First Derm is a store and forward tele-dermatology service that delivers advice for users with skin conditions by a team of international board-certified dermatologists based on two uploaded smartphone images and associated information in text. First Derm gives easy access to a dermatologist for advice, which otherwise can be time-consuming and more expensive for patients. According to the company, about 80 % of their users are helped by over-the-counter medication. 20 % require an additional appointment for further testing and additional treatments.

Over a 10-year period First Derm has received hundreds of thousands of amateur smartphone images from tele-consultations portraying a large number of skin conditions. Using a data set of 63237 images (Inflammatory 44.5k images, Tumor 11k images, Genitalia 4571 images and 3166 Other skin disease images. Where it is estimated that 5-10% are of black skin images. With these images, an AI-algorithm evolved by training a CNN and developed their first classification system version 33.1, with ability to identify 33 diseases in Dermatology and Venereology. This algorithm called Skin Image Search™ (from here on, the AI app) is a free and fast service, available online. The user uploads two image from their smartphone showing their skin condition, and the online service searches its dataset for matching images and within seconds, returns a list with the top 5 most likely diagnoses, in falling order from 1 to 5, hereby referred to as the “top 5”. The AI only analyses the first of the two picture uploaded.

According to the company, the AI app has shown results of 31.6% accuracy in returning the true diagnosis ranked as the most likely. The accuracy in the top 5, e.g. the presence of the true diagnosis among the five proposed differential diagnoses, was 69.9%. The company is continuously experimenting with their CNN and hope to reach higher diagnostic accuracy in future versions with coming updates of the AI app.

### Testing the AI classification system

Collected images were uploaded for automated classification using an online version of the AI app [17] which required uploading two images, one showing the wider body area where the lesion is situated and second close-up photo to allow the classification process to be made as illustrated in figure 1. The top 5 differential diagnoses returned from the AI app against each individual image that was tested were imported into Microsoft Excel sheet. Matching the image’s clinical diagnosis with the returned top 5, each classification was given a score from 1-5 depending on which position the confirmed diagnosis had, score 1 being the most likely. If the confirmed diagnosis was absent in the top 5, the classification was given a score of 0.

**Figure 1:**
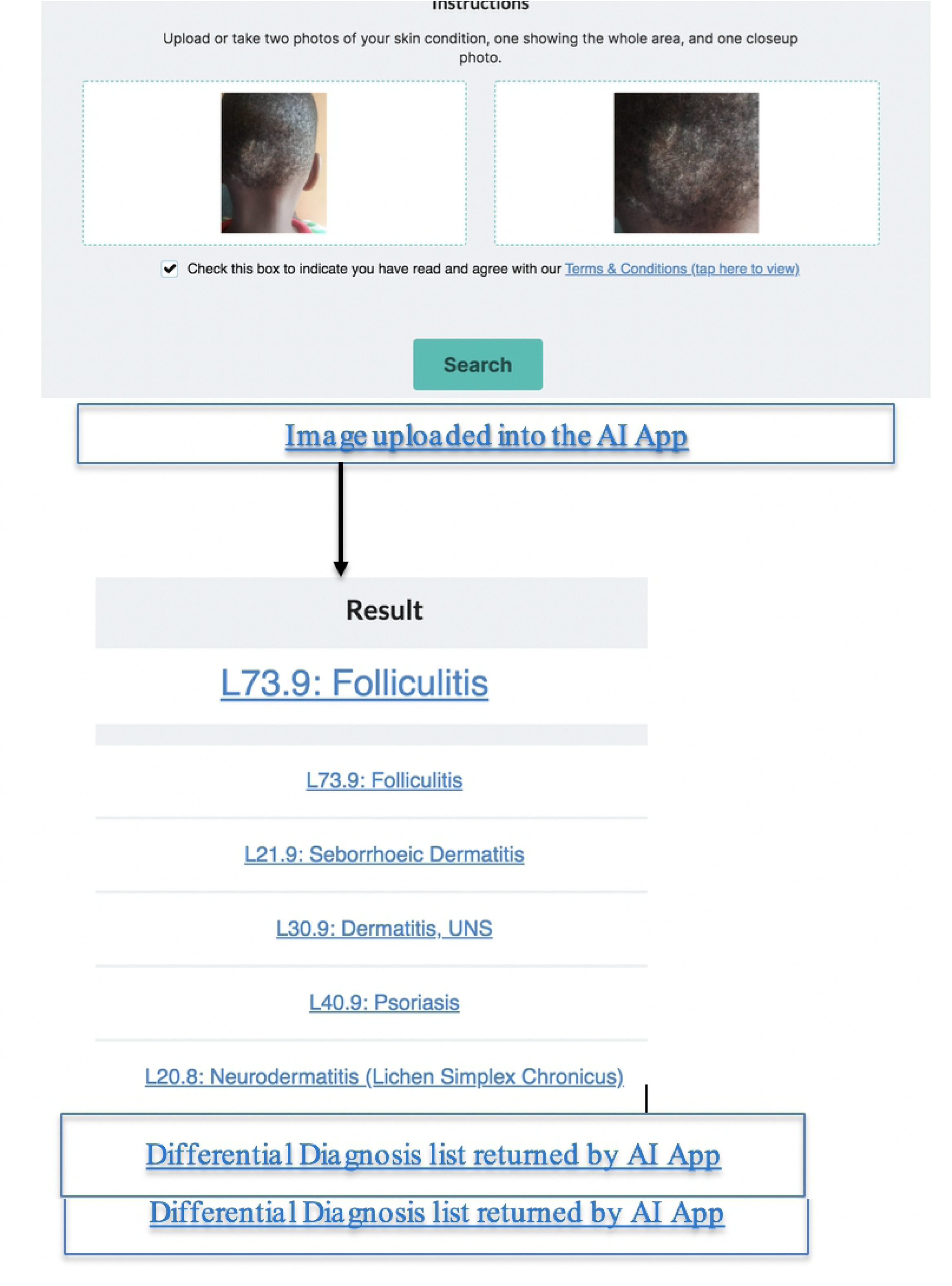
Illustration of the testing process of the AI app.

### Diagnostic accuracy

The overall diagnostic accuracy of the AI app was analyzed as well as the diagnostic accuracy for separate diagnosis, diagnostic groups with a common etiology (e.g. tumors, viral diseases and fungal diseases) or body site (e.g. genital diseases and facial diseases). The dermatological diagnosis analyzed by the AI app including the relevant diagnostic groups are presented in Table 1.

**Table 1:** Dermatological diagnosis with associated diagnostic codes. Presentation of the diagnoses with respective diagnostic codes (ICD-10) for dermatological conditions tested with the AI app in this study.

| Diagnostic group- N (%) | Diseases diagnoses | ICD-10 |
| --- | --- | --- |
| **Bleeding disorders**<br>2 (1.6%) | Ecchymoses | R233 |
| **Burns**<br>4 (3.2%) | Burns | L550 |
| **Chickenpox**<br>4 (3.2%) | Varicella | B01 |
| **Dermatitis**<br>20 (16.2%) | Impetigo<br>Eczema<br>Psoriasis | L01<br>B000<br>L40 |
| **Facial diseases**<br>9 (7.3%) | Acne Vulgaris | L70 |
| **Folliculitis**<br>4 (3.2%) | Folliculitis | L662 |
| **Fungal Diseases**<br>33 (26.8%) | Tinea cruris<br>Tinea corporis<br>Onychomycosis<br>Tinea capitis<br>Athletes foot<br>Oral candidiasis | B356<br>B354<br>B359<br>B350<br>B359<br>B378 |
| **Genital Diseases** |  |  |
| 9 (7.3%) | Pearly Penile Papules<br>Genital Warts<br>Balanitis | L944<br>A630<br>N486 |
| **Insect bite**<br>4 (3.2%) | Insect bite | W570 |
| **Lymphadenitis**<br>2 (1.6%) | Lymphadenitis | L040 |
| **Tumors**<br>18 (15%) | Scar tissue<br>Ganglion cyst<br>Benign Penile papules<br>Corns<br>Confluent and reticulated papillomatosis<br>Hemorrhoids<br>Condylomata acumunata<br>Dermoid cyst<br>Dermatosis papulose Nigra<br>Peri orbital edema<br>Keloid | L905<br>M674<br>L944<br>L84<br>L944<br>1843<br>L944<br>M854<br>L817<br>I890<br>L910 |
| **Skin boils**<br>8 (6%) | Furuncle | L02 |
| **Urticaria**<br>2 (1.6%) | Urticaria | L50 |
| **Wounds**<br>4 (3.2%) | Wounds | B871 |
| **Vitiligo**<br>4 (3.2%) | Vitiligo | L80 |

### Statistical analyses

All data was wrangled and analyzed using R version 3.6.0 on R studio Version 1.2.1335. The data analysis plan was drafted and followed, the data was majorly categorical (92%) and numerical (8%). All categorical variables were nominal apart from the target which was ordinal. The data was cleaned and wrangled using dplyr package and exploratory data analysis using ggplot2 and vcd packages for univariate and multivariate respectively performed on cleaned data and insights derived and communicated.

### Ethics

Institutional approval for usage of the images within TMCG’s electronic medical records was sought. Image file names did not include any patient data and the patients’ identity could not be discerned in the images used in this study. Furthermore, the use of images in this study did not affect patients’ healthcare.

## Results

A total of 173 dermatological images were gathered from TMCG’s electronic medical records 50 images were excluded according to the exclusion criteria above, resulting in a final number of 123 images included and uploaded one by one for classification in the AI app. The gender divide of the study population was 47 males (38 %) and 76 females (62%) with a median age of 23 years. See Table 2 for demographic characteristics of the population. The five most common body sites were genital (20%), trunk (20%), lower limb (14.6%), face (12%) and upper limb (12%). (Table 3).

**Table 2:**
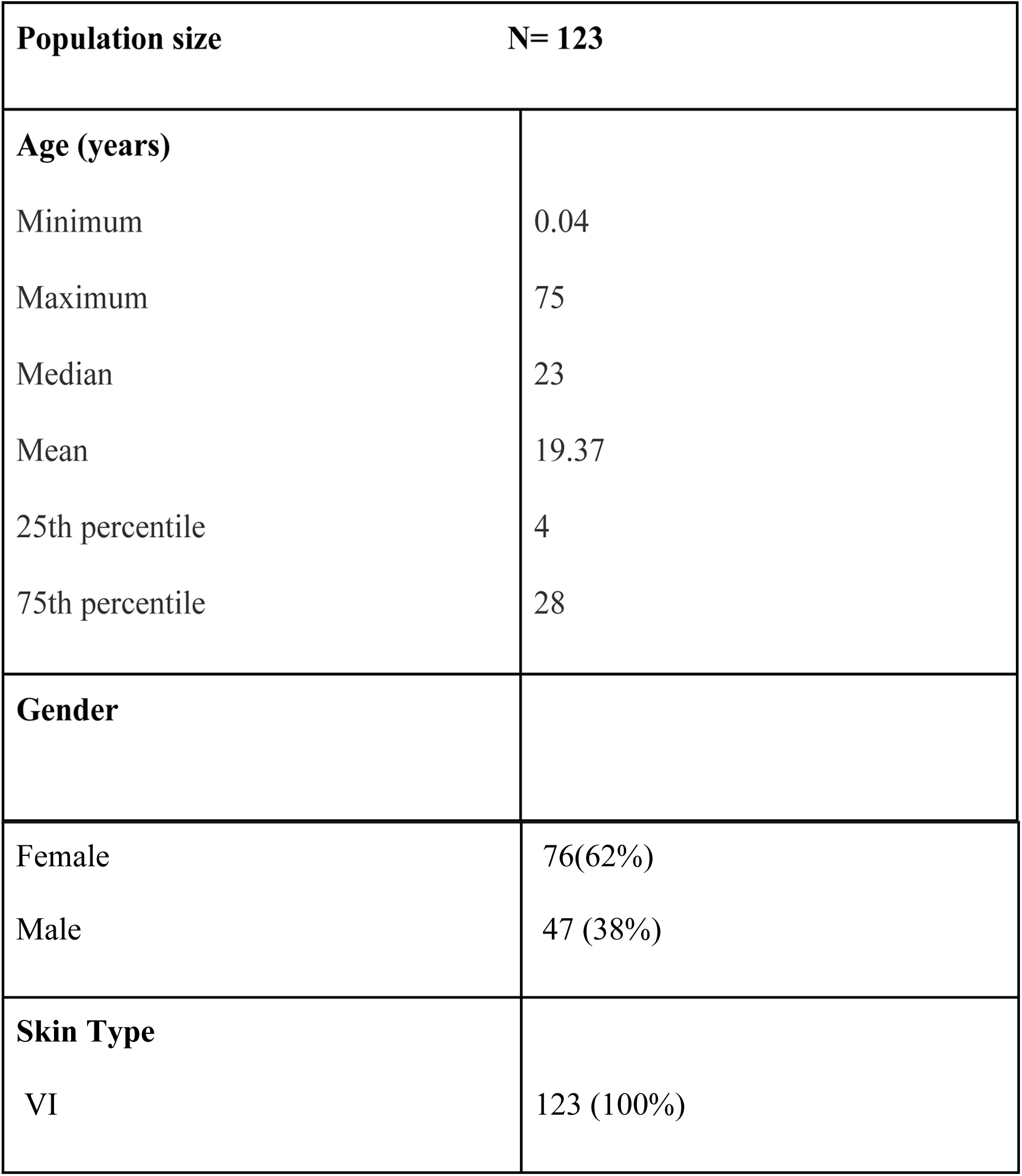
Demographic characteristics.

**Table 3:** Image count in relation to body site.

| **Body Site** | **Image count (%)** |
| --- | --- |
| Anal | 2 (1.6%) |
| Trunk | 1 (0.8%) |
| Ear | 1 (0.8%)) |
| Face | 15 (12%) |
| Genitals | 24 (20%) |
| Lips | 1 (0.8%) |
| Lower limb | 18 (14.6%) |
| Lower limb Extremities | 5 (4%) |
| Neck | 6 (4.9%) |
| Oral Cavity | 6 (4.9) |
| Scalp | 5 (4%) |
| Trunk | 24 (20%) |
| Upper limb | 15 (12%) |

The diagnostic accuracy of the AI app is presented initially with general results i.e. overall disease diagnostic accuracy and thereafter with more specific results i.e. accuracy in relation to other variables i.e. body site and gender.

### Disease diagnostic accuracy of AI

Out of the 123 images uploaded for classification, the AI app was able to place the correct diagnosis among the top 5 in 17% of the images (21/123) and failed to return the correct diagnosis in 83% of the images (102/123). The varying predictability level correctness of the AI app was; 1-8.9%, 2-2.4%, 3-2.4%, 4-1.6% and 5-1.6%; making the true diagnosis (level 1) the largest portion in relation to the other scores in the top 5 (Figure 2). The AI app performed very well on Dermatitis with 80% of uploaded images being predicted with the correct diagnosis (level 1). The AI app performed poorly on Tinea (capitis, corporis and cruris) images yet these had the biggest count.

**Figure 2:**
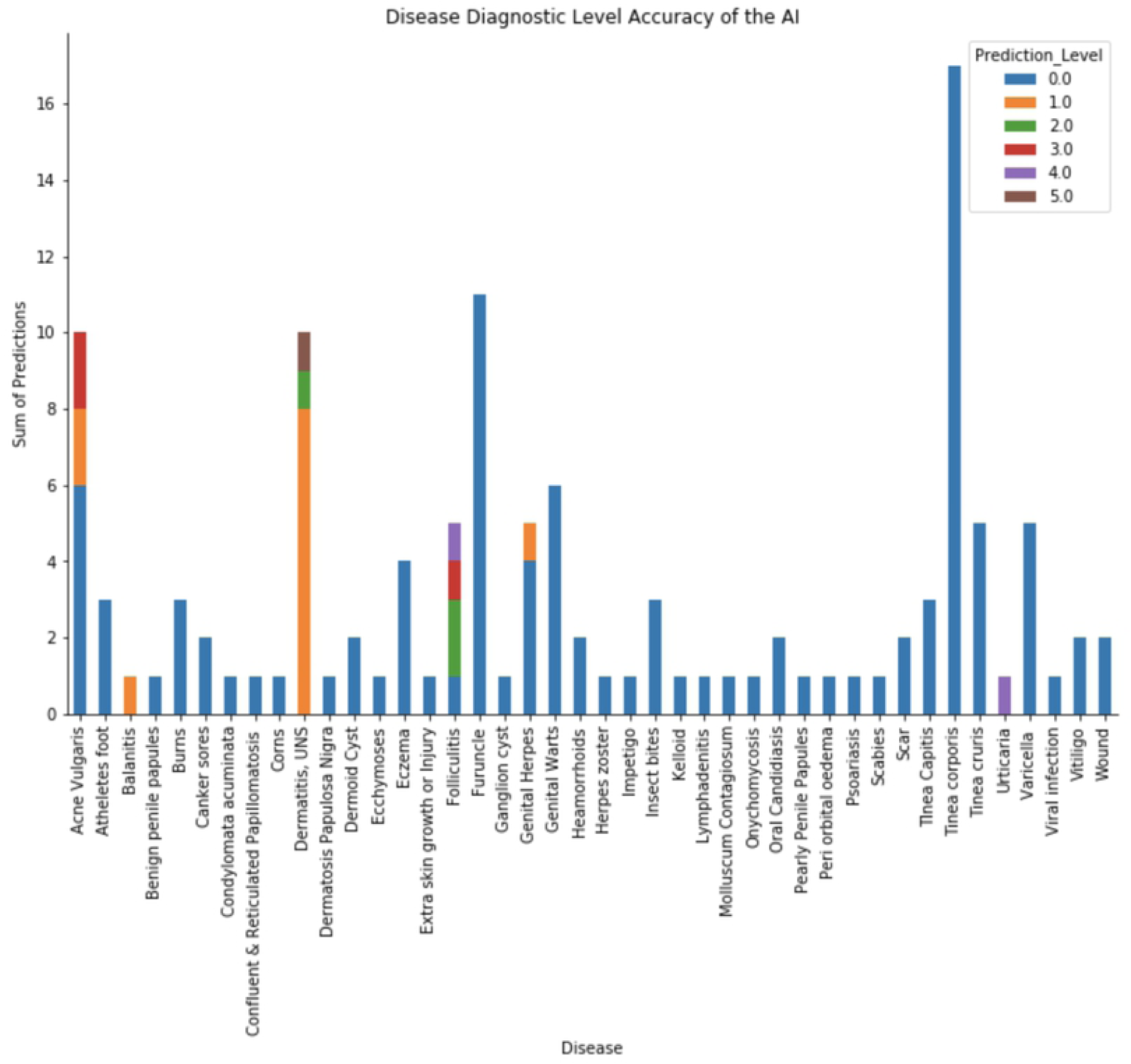
Over all disease diagnostic accuracy of the AI app.

### Body part diagnostic level accuracy of the AI app

Out of the 123 images uploaded; the AI app returned a diagnosis in 62% of all body parts (8/13). The AI app performed well in dermatological images from the face, trunk and genital areas and lowest for lower limb images (Figure 3).

**Figure 3:**
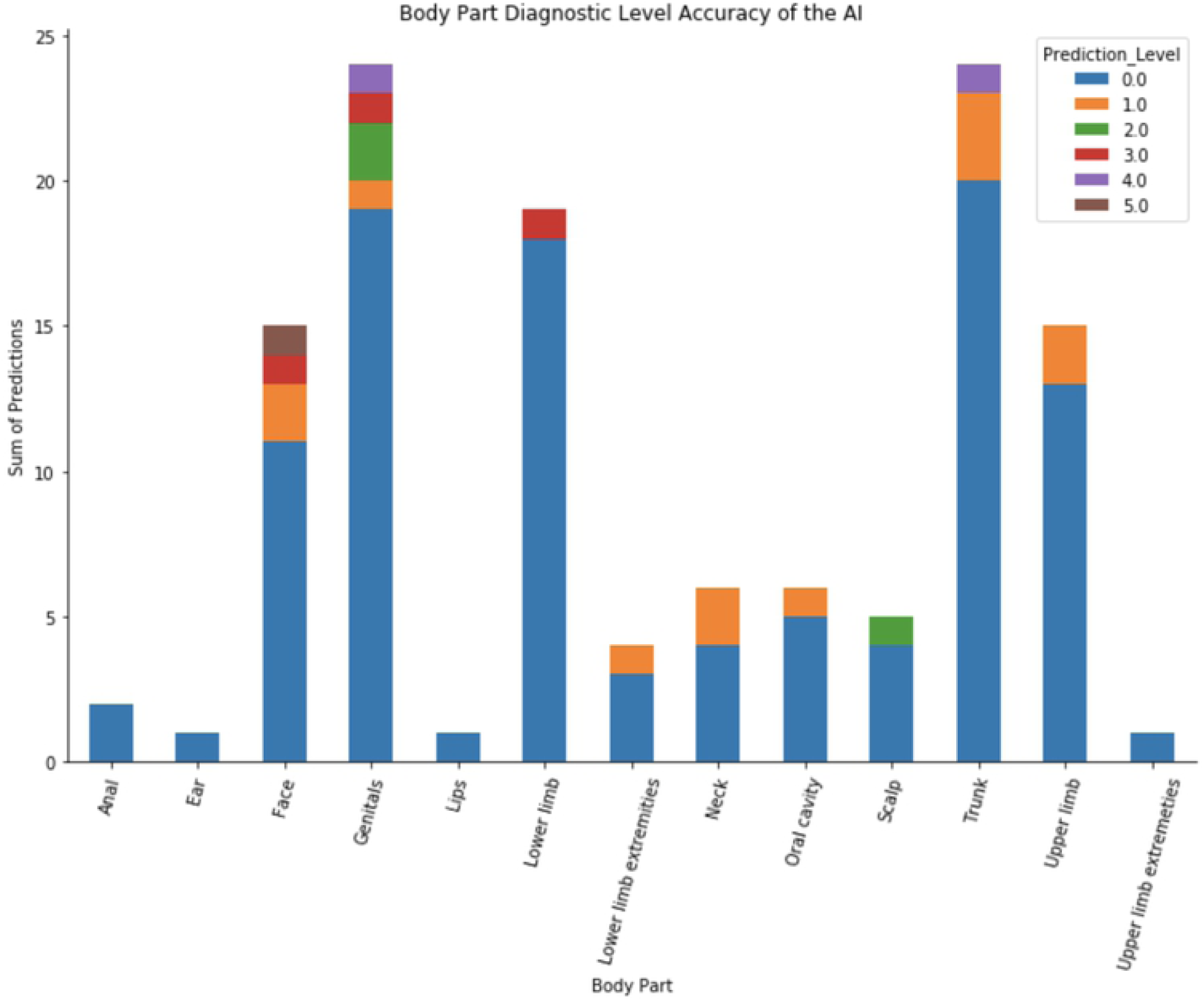
Diagnostic accuracy of the AI app by body part of the dermatological image.

### Clinical diagnosis and gender comparison prediction of the AI app

Overall the AI app performed slightly better among females compared to males for the same dermatological diagnosis. For example; for Dermatitis, the overall performance in females was 70% compared to 20% in males and the same pattern is noted for *Acne Vulgaris* (F-33%, M-11%) and folliculitis (F-60 %, M-20 %). (See figure 4).

**Figure 4:**
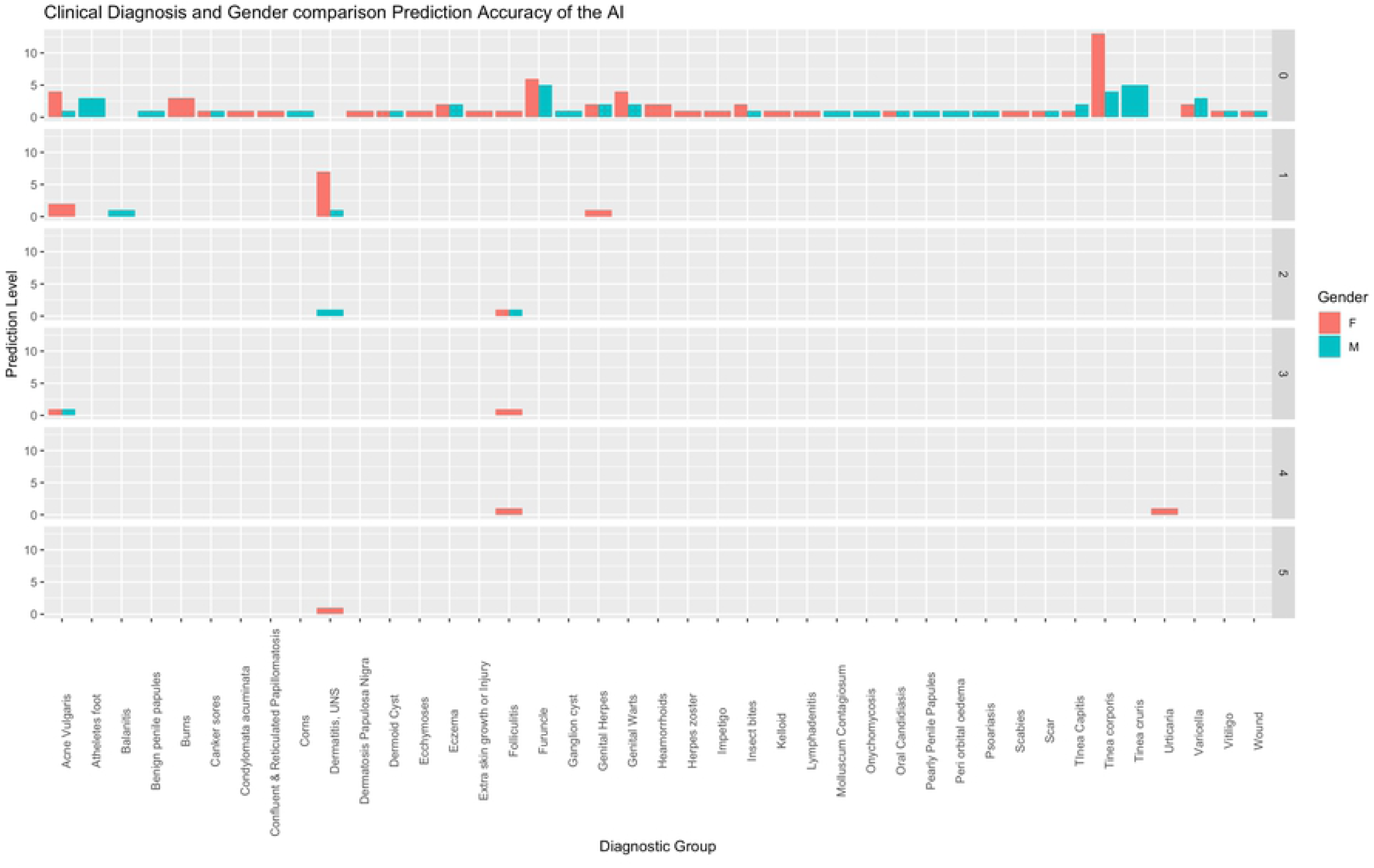
Gender comparison of the diagnostic accuracy of the AI app across dermatological clinical diagnoses.

### Body part and gender comparison prediction of the AI app

AI app predictability of correct diagnosis along body part showed some trends of gender preference with better performance noted in females compared to males. For example; for facial images, the AI app performance among females was 20% compared to 6.7% in males; for genital images, performance among females was 12.4% compared to 8.3% in males; for images of the trunk, performance among females was 8.3% compared to 0% in males etc. (See figure 5).

**Figure 5:**
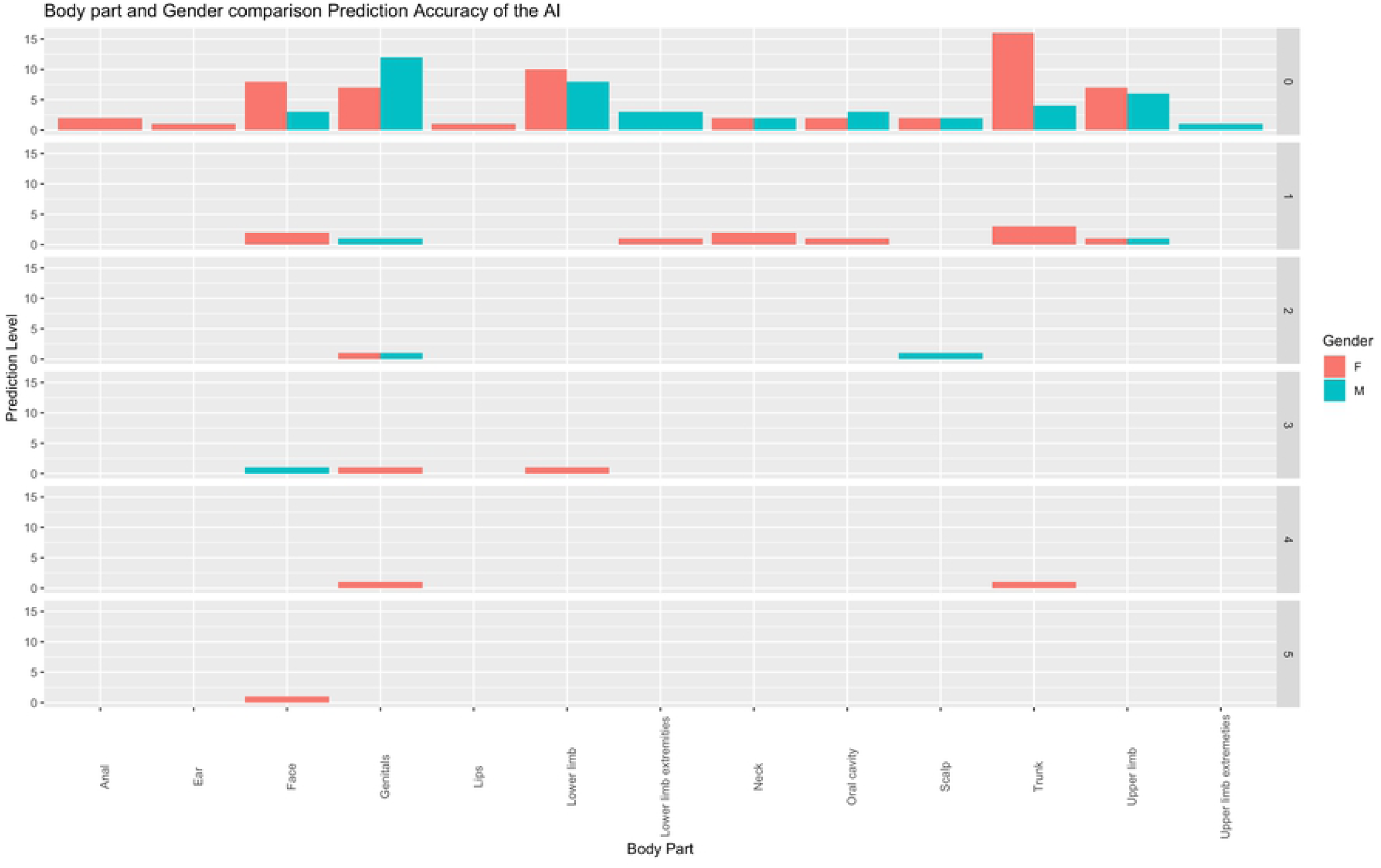
Gender comparison of the diagnostic accuracy of the AI app along body parts of the dermatological images.

## Discussion

First-Derm is an AI-powered dermatology application under development and TMCG a digital health company was chosen to provide an external beta-testing [20] in a black African environment. This manuscript discusses the learning from testing First-Derm with Fitzpatrick 6 skin types (African black skin) dermatological conditions.

Fungal diseases accounted for the majority of dermatological images reviewed (26.8%), these findings relate with the 2010 global burden of disease report which placed fungal diseases as the most prevalent among the top 10 skin diseases globally [4]. Environmental factors including the warm humid weather has been noted to contribute to an increased risk of fungal infestations [21] which are very consistent with the Uganda geographical setting.

The study population was relatively young with a median age of 23 years, this is largely from the fact that these images were from the electronic medical records of a tele-health platform for which studies have always related young age with uptake and usage of digital platforms for medical care [22,23]. This further creates an opportunity to leverage on AI applications for dermatological medical care among the young people who often do not have enough resources and rely on peer to peer consultations when managing skin problems a trend seen with other medical problems like mental, sexual and reproductive health [24].

The overall diagnostic accuracy of the AI app was slightly poor at 17% (21/123) an indicator that First-Derm application was mainly trained based on loads of Fitzpatrick 1&2 skin types (Caucasian skin) and less of Fitzpatrick 5 & 6 skin types (Dark colored skin) who make up the majority of images that were used for this testing. This finding correlates with a report on AI use in voice recognition where it was noted that African voices and accents were being excluded or not being detected [25]. Furthermore, similar observations have been made on facial recognition algorithms which struggle to recognize black faces [26].

Performance of the AI app along specific dermatological skin diagnosis showed interesting results with up to 80% correct diagnosis predicted for Dermatitis and no return (0%) for fungal diseases yet these had the highest count. These findings imply that the AI app was trained adequately on dermatitis datasets compared to fungal images which relates to the fact that the prevalence of dermatitis among western and European settings is high compared to fungal infestations [2,27].

In addition, the AI app performed poorly on tumor diagnostic groups (i.e. scar tissue, dermoid cyst, corn, ganglion cyst). This could partly be explained from the angle that the application had not yet been trained with a number of anatomical structures to clearly delineate out the true body structure on which the skin condition has been taken from. However, the AI app returned a diagnosis in 62% of all body parts (8/13) an indicator that the AI app had been trained with images from a variety of different body sites.

Comparison of the AI app diagnostic accuracy along gender for individual dermatological diagnosis and body part showed a slightly better performance in females compared to males. It is not clear as to what may explain this observation, however, viewing it from the angle of skin tone and texture being smoother among African women than men may explain this observation. The First Derm application like any other AI app is not able to integrate the client’s history to a presenting dermatology inquiry in making its list of differentials, as such instances where history to the occurrence of complaints is critical in making the diagnosis for example in cases of burns and wounds such technology based diagnostic tools are likely to make a miss-diagnosis. This is further echoed by the notion that if an AI program is not trained to perform a particular task it will not be able to execute it [28].

## Conclusions

AI is well suited for classification of skin disease, for any classification to be universal; there is a need for diversity in the images used when training CNNs. Diversity in dermatological image datasets will help reduce biases, the need to include many different kinds of skin complexities; Caucasian, dark-colored, brown and African-black skin colors will help achieve significantly better diagnostic accuracy results. In addition, there is a need to widen the scope of disease conditions trained on the algorithm to include those that may be rare in western/European settings yet common in Sub-Saharan Africa.

## Acknowledgments

We acknowledge the administration at TMCG, whose rich electronic medical records database of images was available for testing. We thank the First Derm team for granting us the opportunity to test their AI application.

